# Genetic modification of primary human B cells generates translationally-relevant models of high-grade lymphoma

**DOI:** 10.1101/618835

**Authors:** Rebecca Caeser, Miriam Di Re, Joanna A Krupka, Jie Gao, Maribel Lara-Chica, João M.L Dias, Susanna L Cooke, Rachel Fenner, Zelvera Usheva, Hendrik Runge, Philip A Beer, Hesham Eldaly, Hyo-Kyung Pak, Chan-Sik Park, George Vassiliou, Brian J.P Huntly, Annalisa Mupo, Rachael JM Bashford-Rogers, Daniel J Hodson

**Affiliations:** Wellcome MRC Cambridge Stem Cell Institute; Department of Haematology, University of Cambridge; Cancer Molecular Diagnostics Laboratory (CMDL), Department of Haematology, University of Cambridge; Wolfson Wohl Cancer Research Centre, Institute of Cancer Sciences, University of Glasgow, Garscube Estate, Glasgow, Scotland, UK; Wellcome Sanger Institute, Wellcome Genome Campus, Hinxton, Cambridge, CB10 1SA, UK; Department of Pathology, Cambridge University Hospitals; Department of Pathology, University of Ulsan College of Medicine, Asan Medical Centre, Seoul, Korea; Wellcome Centre for Human Genetics, Roosevelt Dr, Oxford, OX3 7BN, United Kingdom; Department of Clinical Pathology, Cairo University, Egypt

**Author notes:** **Corresponding Author**: Daniel J Hodson, Wellcome MRC Cambridge Stem Cell Institute, Cambridge Biomedical Campus, Hills Road, Cambridge CB2 0AH, UK.

## Abstract

Sequencing studies of Diffuse Large B Cell Lymphoma (DLBCL) have identified hundreds of recurrently altered genes. However, it remains largely unknown whether and how these mutations may contribute to lymphomagenesis, either individually or in combination. Existing strategies to address this problem predominantly utilize cell lines, which are limited by their initial characteristics and subsequent adaptions to prolonged *in vitro* culture. Here, we describe a novel co-culture system that enables the *ex vivo* expansion and viral transduction of primary human germinal center B cells. The incorporation of CRISPR/Cas9 technology enables high-throughput functional interrogation of genes recurrently mutated in DLBCL. Using a backbone of *BCL2* with either *BCL6* or *MYC* we have identified co-operating oncogenes that promote growth and survival, or even full transformation into synthetically engineered models of DLBCL. The resulting tumors can be expanded and sequentially transplanted *in vivo*, providing a scalable platform to test putative cancer genes and for the creation of mutation-directed, bespoke lymphoma models.

## Introduction

Diffuse Large B Cell Lymphoma (DLBCL) is the most common form of non-Hodgkin lymphoma. Although potentially curable with immunochemotherapy, up to 40% of patients succumb to their disease^1^. In an attempt to unravel the biological basis of DLBCL and to identify new therapeutic opportunities, several groups have recently reported large genomic studies^2–4^. These highlight the considerable genetic heterogeneity of DLBCL and identify hundreds of recurrently mutated genes, copy number alterations and structural variants. Clusters of co-mutated genes suggest the existence of genetic subtypes of DLBCL that may behave differently when exposed to therapeutic agents. Whilst the functional and mechanistic consequences of some of these genetic alterations have been established, for the majority we have little to no understanding of their contribution to lymphomagenesis. To translate these genomic findings into therapeutic progress, it is critical to understand the functional importance and therapeutic relevance of these genetic alterations, both individually and in combination.

Existing model systems used for the functional interrogation of lymphoma genetics consist predominantly of lymphoma cell lines and genetically modified mice. However, both have limitations; cell lines were often established from patients with end-stage, non-nodal or even leukemic phase lymphoma and carry an extensive and biased mutational repertoire, further selected over years or even decades of *in vitro* growth^5–8^. Genetically engineered mice, on the other hand, are costly, time-consuming to generate and therefore unsuitable for high-throughput or combinatorial experiments. Furthermore, the genetic requirements for tumorigenesis in mice do not always accurately reflect those in humans^9,10^. As such, the development of new, preclinical models of lymphoma that can capture its considerable genetic diversity, has been identified as a priority area for lymphoma research^11^.

In common with many of the mature B cell malignancies, DLBCL is thought to arise from the germinal center (GC) stage of B cell differentiation^12,13^. An attractive solution would therefore be to use primary human GC B cells as a platform for *ex vivo* genetic manipulation. Equivalent approaches have proved fruitful for epithelial malignancies^14–17^. However, technical difficulties associated with the *ex vivo* culture and genetic manipulation of human GC B cells, including high manipulation-associated cell toxicity and low transduction efficiency, have obstructed the exploitation of such models to study lymphoma.

Here, we describe an optimized strategy that facilitates proliferation and highly efficient transduction of non-malignant, primary, human GC B cells *ex vivo*. We show that combinations of oncogenes permit long term culture *in vitro*, allowing the system to be used for high-throughput screening of oncogenes and tumor suppressors, and for the creation of genetically customized human lymphoma models that can be studied in immunodeficient mice.

## Results

### Ex vivo growth and transduction of primary human GC B cells

Germinal center B cells are programmed to undergo apoptosis in the absence of survival signals from T follicular helper cells and follicular dendritic cells (FDC). Consistent with this, it is well-established that GC B cells perish rapidly if cultured unsupported *ex vivo*^18^. Previous attempts to support *ex vivo* growth of human GC B cells employed CD40lg transfected fibroblasts in combination with soluble cytokines including IL2, IL4 and IL10^18,19^. Related strategies have used a FDC-like feeder cell termed HK that supported GC survival and allowed short term proliferation when combined with CD40lg^20^. With the increasing appreciation of the importance of IL21 to GC B cell biology^21,22^, later systems have used HK feeder cells combined with CD40lg and IL21^23^. However, proliferation of GC B cells in all these systems was typically limited to a period of up to 10 days^18–20,23^.

We employed a similar system based upon a freshly established culture of modified HK cells, termed YK6 that were immortalized with TERT, P53dd and CDK4 (Figure S1a). These were further engineered to express membrane bound human CD40lg and to secrete soluble IL21, termed YK6-CD40lg-IL21 (Figure S1b). We isolated primary GC B cells (CD38^+^CD20^+^CD19^+^CD10^+^) from pediatric tonsil tissue (Figure 1a), which when grown in co-culture with YK6-CD40lg-IL21 survived and proliferated vigorously for up to 10 days without a requirement for any additional cytokines (Figure 1b&c and Supplementary videos).

**Figure 1.**
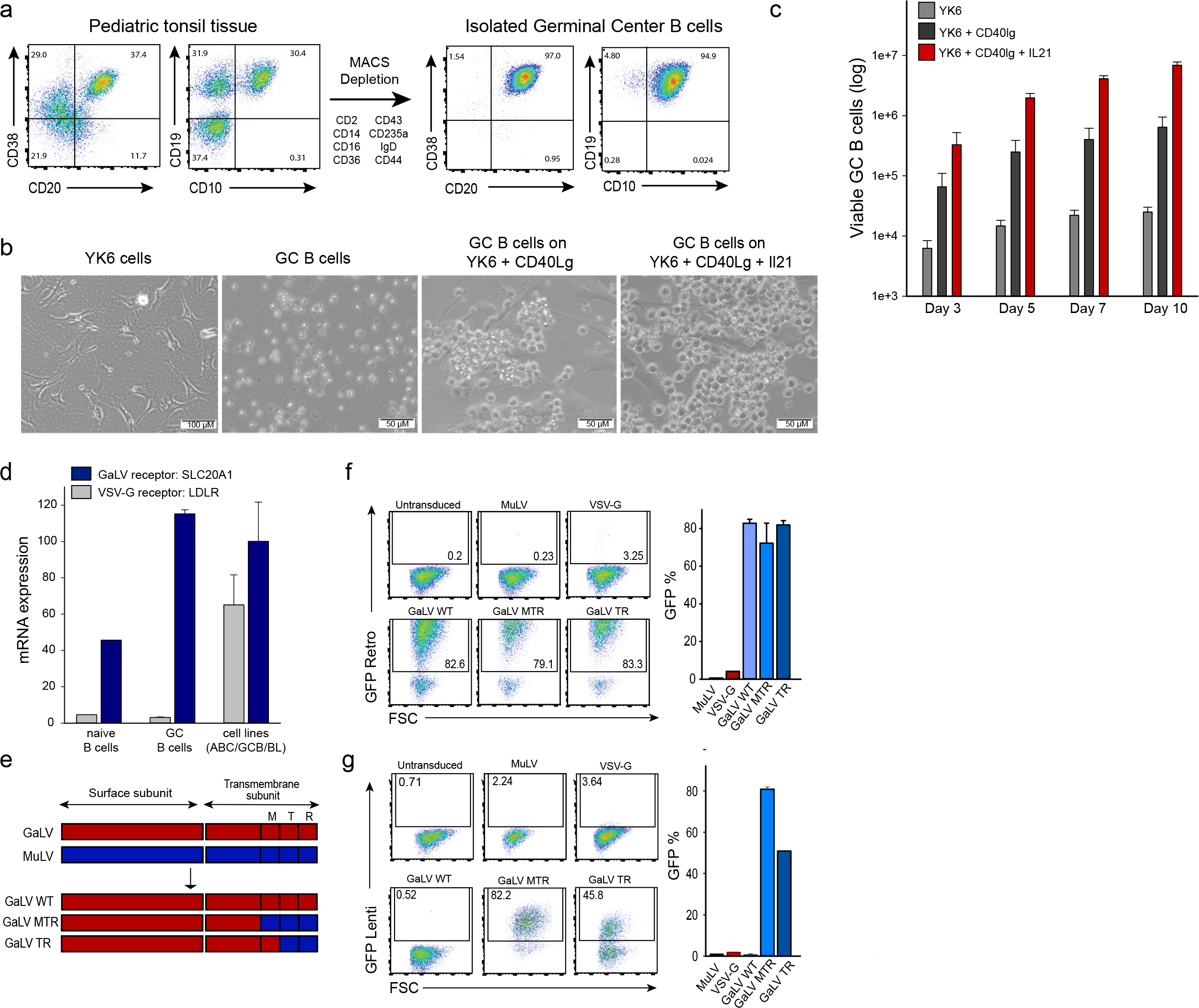
Ex vivo growth and transduction of primary human GC B cells. (a) Representative flow cytometry analysis (*n* > 3) for the expression of GC B cell markers CD38, CD20, CD19 and CD10 in purified GC B cells from pediatric tonsil tissue. The strategy for negative selection of GC B cells is shown. (b) Representative images are shown of YK6 cells, GC B cells alone or GC B cells cultured on either YK6-CD40lg or YK6-CD40lg-IL21 feeder cells. Scale bar represents 50μm or 100μm as indicated. (c) Primary human GC B cells were cultured with YK6 control, YK6-CD40lg or YK6-CD40lg-IL21 feeder cells. Illustrated is bar graph showing the number of viable cells (± Standard Error, n=5) over four timepoints. Viable cells were determined by flow cytometry and counting beads. (d) Bar graph showing the relative expression of *SLC20A1* (GaLV receptor) and *LDLR* (VSV-G receptor) in naïve (n=1), GC B cells (n=3) and ABC/GCB DLBCL and Burkitt cell lines (n=6) as analyzed by RNA-seq. mRNA expression values were calculated as counts per million reads (CPM). Error bars indicate ±Standard Error. (e) Schematic of the retroviral and lentiviral MuLV-GaLV fusion envelopes, GaLV_WT, GaLV_MTR and GaLV_TR. M = transmembrane region, T = cytoplasmic tail, R = R peptide, SU = surface subunit, TM = transmembrane subunit^28^. (f&g) Primary human GC B cells were transduced with a retroviral control (f) or lentiviral control (g) construct using GaLV-MuLV fusion envelope constructs as well as VSV-G and MuLV. Three days after transduction, transduction efficiencies in primary human GC B cells were determined by expression of GFP. Error bars indicate ±Standard Deviation, n=3. FSC, forward scatter.

In line with previous observations in human B cells^24,25^ we were unable to transduce human GC B cells with amphotrophic or VSV-G pseudotyped virus. Peripheral blood B cells have previously been transduced using virus pseudotyped with a Gibbon Ape Leukemia Virus (GaLV) envelope^26^, the receptor for which is *SLC20A1*^27^. RNA-Seq showed that human GC B cells express high levels of *SLC20A1*, but very low levels of the VSV-G receptor *LDLR* (Figure 1d). Thus, we proceeded to test the GaLV viral envelope to transduce primary GC B cells. To permit lentiviral transduction, we generated a series of GaLV-MuLV fusion constructs based on previous reports^26,28^ (Figure 1e) and identified a fusion construct that permitted high efficiency transduction with both retroviral (Figure 1f) and lentiviral (Figure 1g) constructs of human primary GC B cells cultured on YK6-CD40lg-IL21 feeders. Interestingly, the GaLV envelopes also enabled the transduction of primary human DLBCL cells supported on YK6-CD40lg-IL21 cells (Figure S1c).

### Long term expansion of human germinal center B cells ex vivo

We proceeded to use this culture-transduction system to introduce into human GC B cells oncogenes that are commonly deregulated in human lymphoma. Out of five genes tested, no single gene was able to prolong the survival of primary GC B cells cultured in our system (Figure 2a&b). However, *BCL2* when co-expressed with either *MYC* or *BCL6* overexpression did lead to long term expansion and survival of transduced GC B cells in culture. These cells continued to expand and proliferate vigorously in culture beyond 100 days. We also tested other transcription factors associated with the germinal center reaction, and their lymphoma-associated mutants, in combination with BCL2 in a pooled, competitive culture. This showed initial expansion of cells transduced with *MEF2B* Y69H, a mutation commonly found in DLBCL and follicular lymphoma^29^. However, by day 59, cultures were dominated by *BCL6*-transduced cells suggesting this as the transcription factor best able to promote long-term growth of GC B cells *ex vivo* (Figure 2c, Supplementary Table 1). Flow cytometry after 10 weeks of culture showed that cells transduced with *BCL2* and *BCL6* maintained expression of surface markers reminiscent of GC B cells including CD19, CD20, CD22, CD38, CD80 and CD95 (Figure 2d). Cells expressed both CD86 and CXCR4 markers, an immunophenotype intermediate between light and dark zone GC B cells (Figure 2d). Cells transduced with *BCL2* and *MYC* remained viable and proliferated but downregulated CD20 and CD19, consistent with differentiation towards plasmablasts (Figure S1d). The plasma cell marker CD138 was not expressed by either *BCL2*/*MYC* or *BCL2*/*BCL6* transduced cells (Figure S1e). We compared gene expression profiles of freshly isolated and transduced GC B cells cultured *ex vivo* at early (5 days) and late (10 weeks) time points (Figure 2e). As anticipated, this showed enrichment of a STAT3 signature in cultured cells consistent with ongoing IL21 stimulation. While freshly isolated GC B cells were enriched for expression of centroblast genes, the cultured and transduced cells adopted a gene expression profile more similar to that of centrocytes, consistent with ongoing CD40 stimulation. Importantly, the centrocyte is the stage of GC differentiation most similar to DLBCL^30^. Transcriptome analysis was also compared with that of six cell lines commonly used as models of GC-derived lymphomas, including the main subtypes of DLBCL and Burkitt lymphoma. When compared to a signature of germinal centre expressed genes (GCB-1)^31^, long-term *BCL6*-transduced cells clustered more closely with GC B cells than did the cell lines (Figure 2f & Figure S1f).

**Figure 2.**
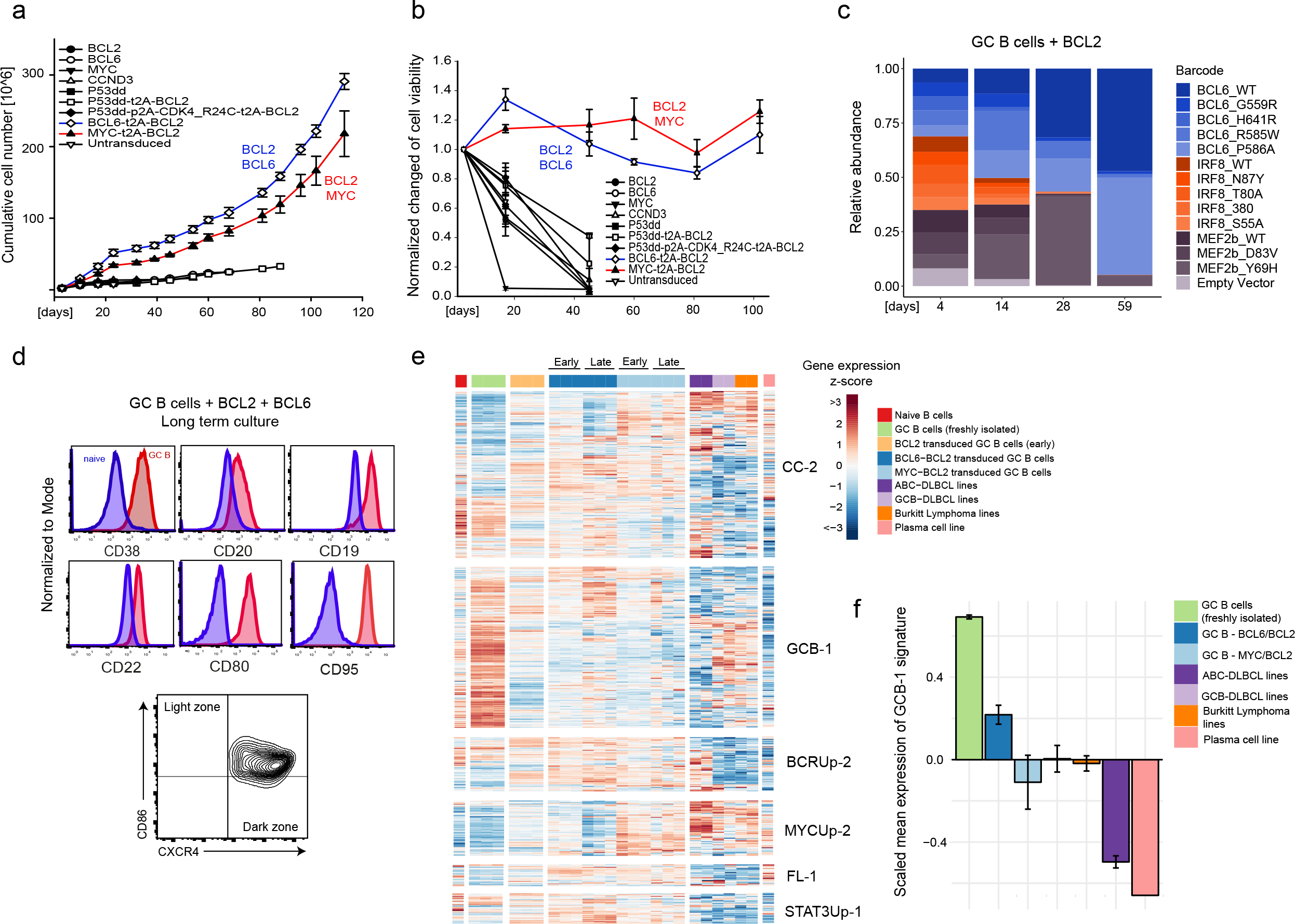
Long term expansion of human germinal center B cells ex vivo. (a) Primary human GC B cells were transduced with the indicated oncogenes and oncogene combinations and cultured separately for up to 120 days. Graph shows calculated theoretical absolute cell numbers (± Standard Error, n=3). Viable cells were assessed by trypan blue exclusion. (b) Primary human GC B cells were transduced with different oncogenes and oncogene combinations and monitored by flow cytometry. Graph shows the change in cell viability assessed by scatter characteristic by flow cytometry (± Standard Error, n=3). (c) Primary human GC B cells were transduced with *BCL2* in combination with other transcription factors in a pooled, competitive culture. Graph shows relative abundance of transcription factors or their mutant versions over 4 different timepoints (n=3). (d) Primary human GC B cells were transduced with the oncogenic cocktail *BCL2* and *BCL6* and cultured to day 73. Representative flow cytometry analysis (*n* = 3) for the expression of the GC B cell markers CD38, CD20, CD19, CD80, CD22, CD95, CXCR4 and CD86. Red histograms show GC B cells compared to primary human naïve B cells (blue). (e) Heat map of gene expression of freshly isolated GC B cells (n=3), transduced GC B cells (*BCL2-BCL6, BCL2-MYC*) cultured *ex vivo* for 5 or 73 days (n=3), Plasma cell line (n=1), naïve B cells (n=1) and lymphoma cell lines (TMD8, HBL1, SUDHL4, DOHH2, Mutu and Raji, n=6). Illustrated are the selected gene expression signatures - CC-2, GCB-1 as well as BCRUp-2, MYCUp-2, FL-1 and STAT3Up-1. Order of genes in the signatures was determined by hierarchical clustering. (f) Bar chart showing the scaled mean expression of the germinal center B cell signature, GCB-1 in freshly isolated GC B cells (n=3), transduced GC B cells (*BCL2-BCL6, BCL2-MYC*) cultured *ex vivo* for 73 days (n=3), Plasma cell line (n=1) and lymphoma cell lines (TMD8, HBL1, SUDHL4, DOHH2, Mutu and Raji, n=6). Error bars represent the standard error of the mean from independent biological replicates.

Overall, these results suggest that transduced primary human germinal center B cells can be cultured long-term *ex vivo*, retaining characteristics of the initial GC B cell that are shared with DLBCL cells. This represents a valuable new model system for the functional interrogation of genes involved in germinal center lymphomagenesis.

### High-throughput screening for tumor suppressor genes in cultured primary human GC B cells

We wished to use the system for the high-throughput study of putative tumor suppressor genes (TSGs) in lymphoma. We hypothesized that many tumor suppressor pathways are already inactivated in lymphoma cell lines, and as such, primary GC B cells should be more sensitive in identifying a competitive growth or survival advantage following TSG inactivation. Robust expression of Cas9 was achieved using a stable Cas9 retroviral packaging line (Figure S2a) and initial experiments confirmed efficient gRNA-directed targeting in primary, human, GC B cells *ex vivo* (Figure S2b & c). We therefore created a lymphoma-focused CRISPR gRNA library composed of 6000 gRNAs targeting a total of 692 genes reported to be mutated or deleted in human lymphoma, along with 250 non-targeting control guides. Each gene was targeted by up to 9 gRNAs (Figure S2d) and deep sequencing revealed that 99% of gRNAs were within four times of the mean in frequency (Figure 3a). The library was transduced into primary GC cells shortly after their transduction with *BCL2*, *BCL6* and *Cas9* cDNAs. Figure 3b shows an experimental scheme of the CRISPR screening. Cas9 and gRNA constructs were marked with fluorescent proteins to allow selection to be visualized by FACS. Whilst Cas9 and gRNA dual infected cells comprised only 10% of all cells at day 4 this population expanded to 90% by day 88 of culture (Figure S2e), suggesting strong selection for one or more of the library gRNAs. Genomic DNA was sequenced at intervals and a CRISPR gene score was generated for each gene (Figure 3b).

**Figure 3.**
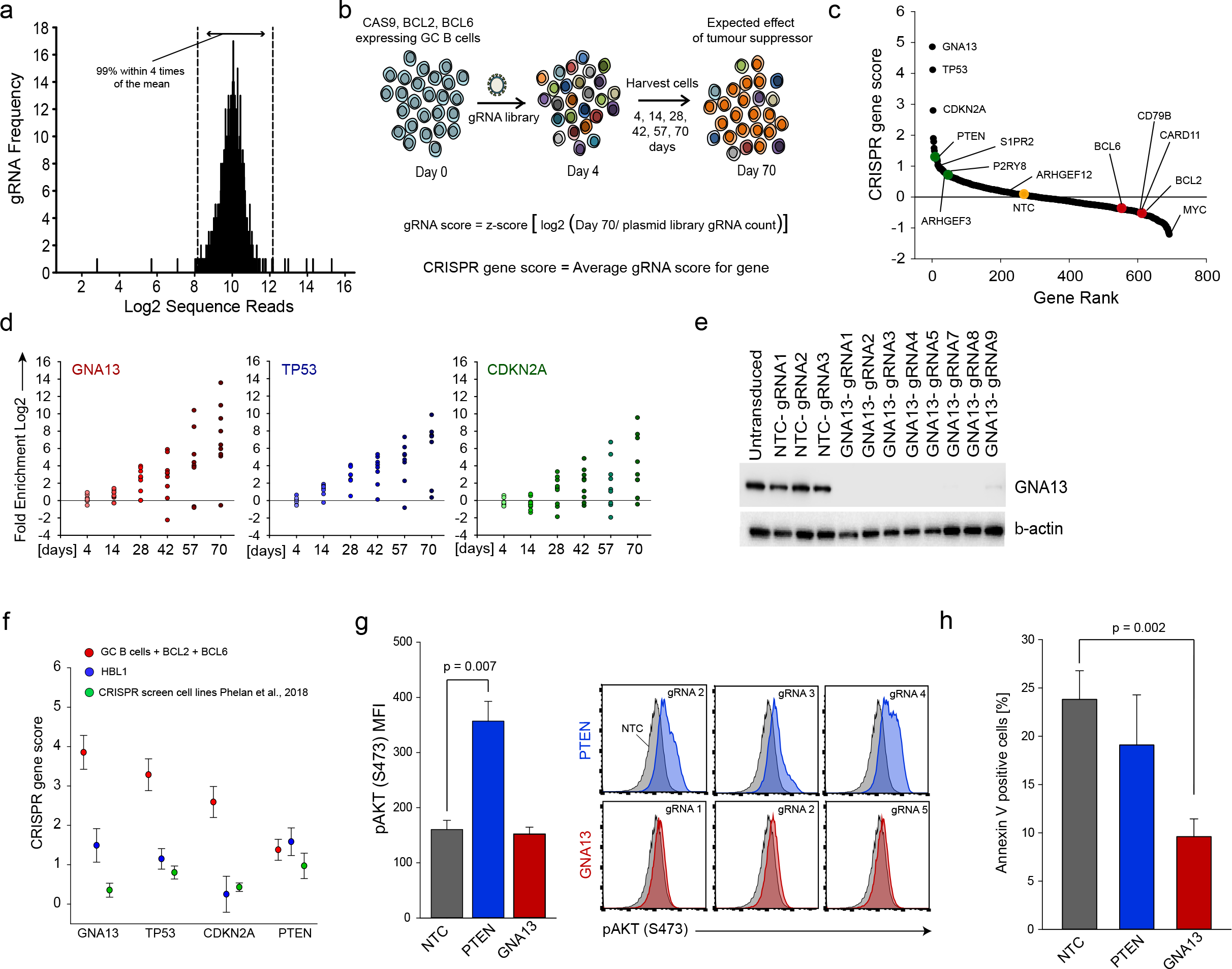
High-throughput screening for tumor suppressor genes in cultured primary human GC B cells. (a) Illumina sequencing of the lymphoma-focused CRISPR library revealed that 99% of sequence reads were within 4 times of the mean. Illustrated is the gRNA frequency vs log2 sequence reads. (b) Outline of experimental design and mathematical formulas used. (c) Rank-ordered depiction of CRISPR gene scores at Day 70 (log2 scale) from highest to lowest. Selected tumor suppressor genes as well as oncogenes are highlighted in green and red, respectively. Data points above the horizontal line are positively enriched. CRISPR gene score for non-targeting control (NTC) (n=250) is 0.22. Representative of 3 experiments. (d) Illustrated is log2 fold enrichment for *GNA13*, *TP53* and *CDKN2A* normalized to non-targeting control gRNAs at the indicated timepoints after transduction with the CRISPR library. Circles represent individual gRNAs for the indicated gene. Data points above the horizontal line are positively enriched. Representative of 3 experiments in cells from independent donors. (e) The ABC-DLBCL cell line HBL1-Cas9 was transduced with 8 gRNAs against *GNA13* and 4 against non-targeting control. Cells were harvested 10 days after transduction and a western blot performed to validate knock-down. β-actin was used as a loading control. Representative of > 3 experiments. (f) CRISPR gene score for *GNA13, TP53, CDKN2A* and *PTEN* in GC B cells transduced with *BCL2* + *BCL6* (n=3), HBL1 (n=1) and compared to published^42^ CRISPR gene scores in lymphoma cell lines (n=11). Error bars indicate ± Standard Error of all gRNAs targeting the indicated gene. (g) pAKT levels following *GNA13* and *PTEN* deletion in primary human GC B cells was monitored by intracellular FACS staining for pAKT (S473). A representative example is shown of 3 PTEN gRNAs (blue)/ *GNA13* gRNAs (red) against 3 non-targeting control (NTC) gRNAs (grey). Bar chart illustrates the mean fluorescence intensity of all gRNAs (*GNA13*=6, *PTEN*=4, NTC=3) for the indicated gene (± Standard error). The p value was calculated from t test. (h) Cell survival following *GNA13* and *PTEN* deletion in primary human GC B cells was monitored by annexin-V and 7-aminoactinomycin D (7AAD) staining and analyzed by flow cytometry. Bar chart illustrates Annexin-V positive cells. The p value was calculated by t test.

Genes that showed the greatest enrichment during culture over 10 weeks included well-established tumor suppressors such as *TP53*, *CDKN2A* and *PTEN* (Figure 3c), thus validating the ability of our system to detect *bona fide* TSGs. Interestingly, the greatest enrichment was seen for *GNA13* (Figure 3c), which encodes the G protein subunit α13. Inactivating mutations of *GNA13* are common in DLBCL and BL^32,33^ but rare in other forms of cancer, where, in contrast, amplification may be more common (Figure S2f)^34,35^. As such, *GNA13* can be considered as a germinal center specific TSG. Enrichment was seen for 8 out of 9 gRNAs targeting *GNA13* over different timepoints, with similar results seen for *TP53* and *CDKN2A* (Figure 3d), and was reproduced in replicate screens performed using GC B cells from three separate donors (Supplementary Table 2). All *GNA13* gRNAs led to effective depletion of GNA13 (Figure 3e), apart from one which was associated with presumed off-target toxicity and further confirmed in a cell line (Figure S2g). We performed a parallel screen using the lymphoma cell line HBL1 (Figure S2h) and also compared data from recent published CRISPR screens (Figure 3f). In these cell line experiments, enrichment of gRNAs targeting TSGs was much more modest. This highlights a unique strength of this system to identify genetic changes associated with enhanced growth and survival; a phenotype that is hard to identify using heavily mutated cell lines, already optimized for *in vitro* growth.

GNA13 acts downstream of the G-protein coupled receptors S1PR2 and P2RY8 and enrichment for both genes was observed in our screens (Figure 3c). Mouse knockout studies have suggested that suppressed activity of this pathway in lymphoma may allow egress from the germinal centre and increase cell survival secondary to enhanced AKT activity^33,36,37^. In contrast, other studies suggest a pro-survival effect in DLBCL that is independent of AKT activity^38^. We therefore quantified pAKT levels in *ex vivo* GC B cells transduced with gRNAs against *GNA13*, *PTEN* or non-targeting controls, and stimulated on YK6-CD40lg-IL21 feeder cells (Figure 3g). Although pAKT was increased in *PTEN*-depleted cells, no increase was seen in *GNA13*-depleted cells. However, *GNA13* depletion did lead to a marked reduction in apoptosis in cultured primary GC cells (Figure 3h), but no change in cell proliferation (Figure S2i). This confirms AKT-independent, enhanced cell survival as the explanation for the competitive advantage seen following *GNA13* depletion in this culture system.

These experiments highlight how primary *ex vivo* GC cells can be used both for high-throughput as well as gene-focused functional experiments. They demonstrate how the system is especially suited to the identification of genetic alterations associated with increased competitive fitness, a phenotype that is hard to induce in established cell lines. Finally, the ability to identify lymphoma-specific TSGs strongly supports the validity of this system for the study of lymphoma genetics.

### Transduced human GC B cells form tumors in immunodeficient mice that recapitulate the features of human high-grade B cell lymphoma

To examine the ability of the culture-transduction system to recapitulate lymphomagenesis *in vivo*, we transduced primary, human GC B cells with combinations of oncogenic alterations commonly found in DLBCL and injected them in Matrigel into immunodeficient mice (Figure 4a). Although sufficient for long-term, feeder-dependent growth *in vitro*, transduction with *BCL2* and *BCL6*, with or without the addition of a dominant negative *TP53* (P53dd)^39^ was insufficient for tumor formation *in vivo*. However, the addition of a fourth oncogene (*BCL6, BCL2, P53dd* & *CCND3*) led to tumor formation with a median of 112 days (Figure 4a). The combination of *MYC*, *BCL2* & P53dd, led to tumor formation with a median of 111 days and the combination of *MYC, BCL2* & *BCL6* resulted in tumor formation with a median of 108 days. The most potent combination tested; *MYC*, *BCL2*, P53dd and *CCND3*, resulted in tumor formation in all mice within 38 days. Notably, tumors engrafted with a 100% penetrance and could be derived from multiple donors, excluding the possibility of donor-derived occult mutations contributing to transformation. Flow cytometry showed cells to be strongly positive for markers of all transduced oncogenes, suggesting potent selection during tumorigenesis (Figure 4b).

**Figure 4.**
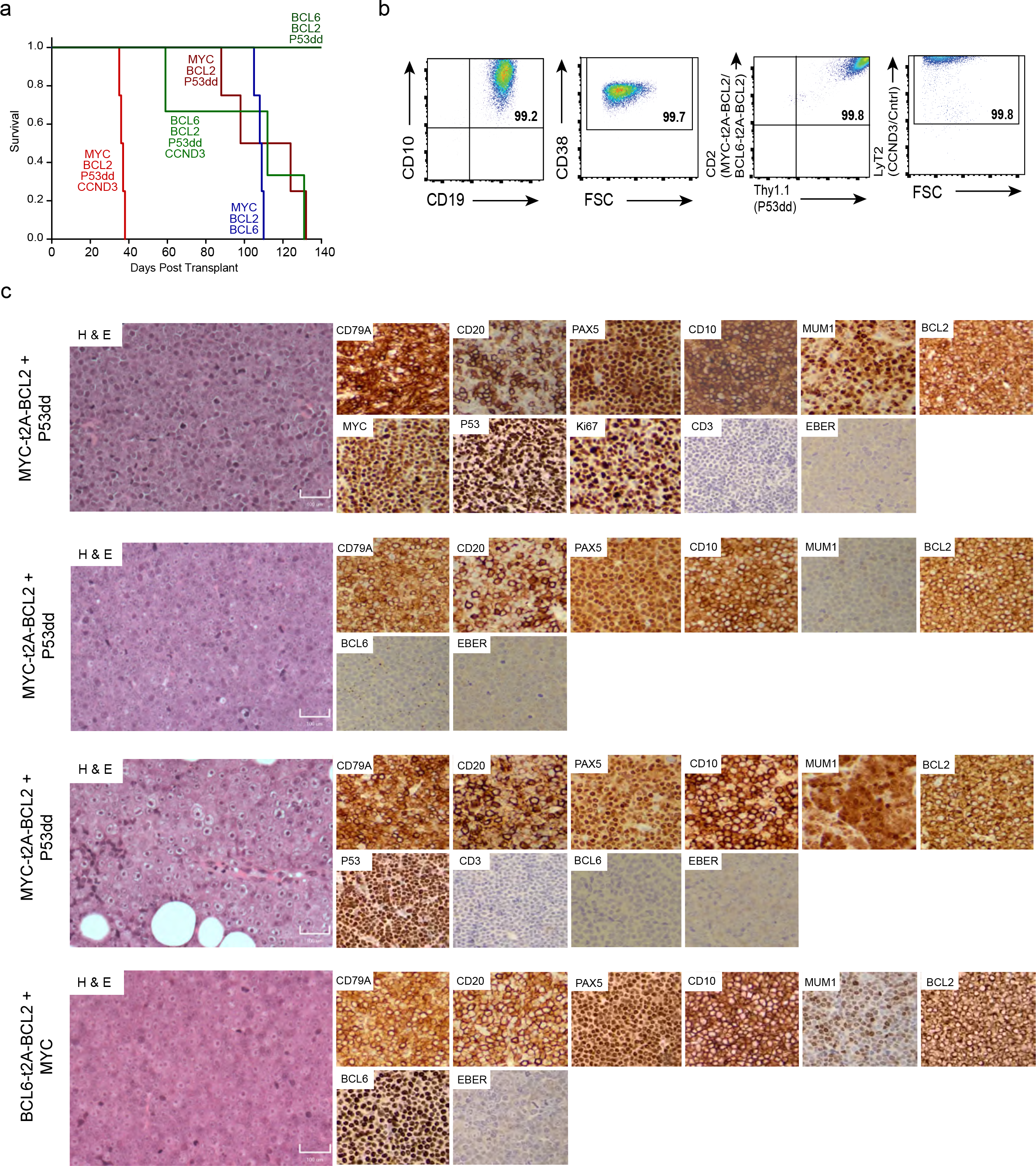
Transduced human GC B cells form tumors in immunodeficient mice that recapitulate the features of human high-grade B cell lymphoma. (a) Primary human GC B cells (as well as YK6-CD40lg-IL21 cells) transduced with the indicated oncogene cocktails were injected subcutaneously into NOD/SCID/gamma mice (n = 3-4 per cohort) and monitored for palpable tumors. Mice were culled when tumors reached 12mm in size. Survival of the recipient mice is plotted as a Kaplan-Meier curve. (b) Cells isolated from tumors were stained for the viral transduction markers CD2 (*MYC-t2A-BCL2* or *BCL6-t2A-BCL2*), Thy1.1 (P53dd), LyT2 (*CCND3*/Cntrl) and B cell markers CD19, CD10 and CD38 and were analyzed by flow cytometry. A representative example is shown. FSC, forward scatter. (c) H&E and Immunohistochemistry images for the indicated proteins are shown (Magnification 20×). Scale bar, 100μM. Four representative tumors from mice described in (a) are shown.

Histological examination revealed diffuse sheets of medium to large, atypical lymphoid cells with frequent mitoses, closely mimicking the appearances of human high-grade B cell lymphoma (Figure 4c & Figure S3 & Figure S4). Immunoblastic and Burkitt-like appearances were seen in some tumors. Immunohistochemistry showed expression of the B cell markers CD19, CD20, CD79A and PAX5 in the majority of tumors (Figure 4c & Figure S3 & Figure S4). In contrast to our *in vitro* observations where cells transduced with *MYC* but not *BCL6* downregulated expression of CD20, most *MYC*-driven tumors expressed strong surface CD20. The germinal center marker CD10 was expressed in approximately half of tumors (Figure 4c & Figure S3). Importantly, all tumors were negative for EBER (ISH) confirming that latent EBV genes did not contribute to lymphomagenesis in these tumors. Harvested tumor cells could be expanded *in vitro* and serially retransplanted back into immunodeficient mice (Figure S5a), thus functioning as a robust laboratory model system.

To establish the clonality of tumors, we performed deep sequencing of PCR amplified immunoglobulin heavy chain variable gene regions to assess the percentage of unique BCR sequences in each sample. This revealed that clonality was increased in primary tumor samples compared to the original donor cells and was also increased in retransplants compared to primary tumors (Figure 5a). BCR network plots showed that tumors with four oncogenic hits were polyclonal (Figure 5b). In contrast, cells transduced with just three oncogenic hits, which formed tumors with a longer latency, were oligoclonal (Figure 5b). This suggests that the combination of 4 oncogenic events (*MYC, BCL2, CCND3* & *P53*) is by itself sufficient for transformation of human GC B cell. In contrast, further oncogenic events are required for lymphomagenesis in cells transduced with just three of the above constructs. To identify these co-operating oncogenic events, we performed targeted sequencing using a hematological malignancy panel of 292 genes. A subclonal *NRAS* G13A mutation (VAF 0.03) was detected in the oligoclonal tumor arising from *MYC, BCL2*, P53dd transduced cells (Figure 5c). This mutation became clonal when retransplanted into secondary recipients confirming its role in the pathogenesis of those tumors (Figure 5c). Mutation at this codon has been reported previously in DLBCL^32^ as have other activating mutations of NRAS^2^. We observed copy number increase for the experimentally transduced gene *BCL6* (Figure S5b) but saw no evidence of any significant aneuploidy in any tumor (Figure S5c). In the polyclonal tumors, subclonal mutations with VAF <0.05 were detected in several genes commonly mutated in DLBCL including a frameshift variant in *S1PR2* and missense mutations in *GNA13, NOTCH2, CREBBP, EP300, SOCS1* and *BCL6* (Figure 5c) (Supplementary Table 3). The significance of these mutations to tumor formation is uncertain, however some of these genes are typical targets of aberrant somatic hypermutation suggesting the possibility of ongoing somatic hypermutation in these lymphomas. To investigate this possibility, we analyzed the variable region sequence of dominant clones detected in the IgH clonality assay. As expected, given their germinal center origin, almost all clones showed evidence of diversification from the germline V gene sequence (Figure 5d&e). Importantly however, clones also showed evidence of ongoing diversification of the hypervariable regions (Figure S5d). This suggests that AID mediated somatic hypermutation remained active during the process of tumor formation.

**Figure 5.**
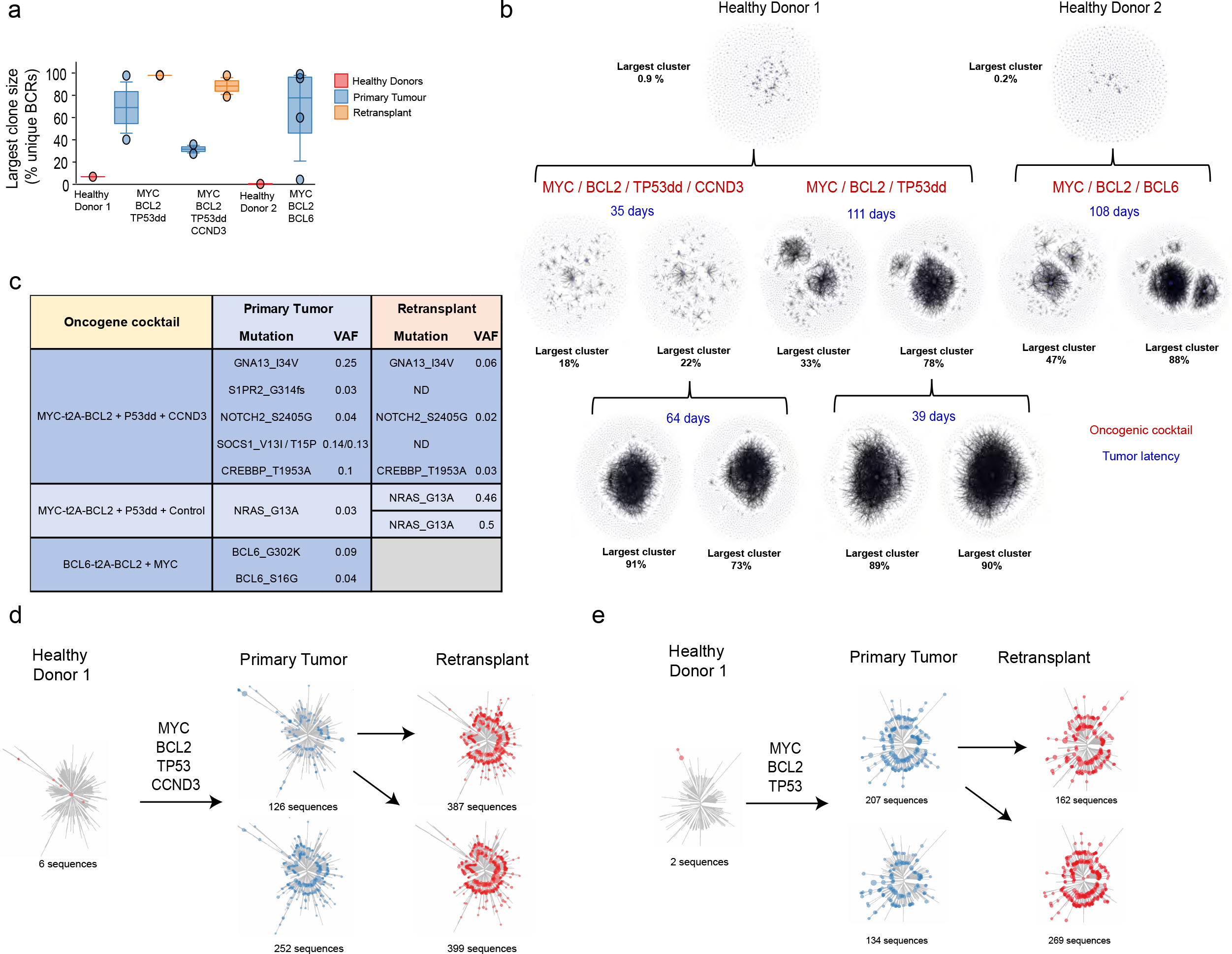
Transduced human GC B cells form tumors in immunodeficient mice with ongoing somatic hypermutation. (a) Boxplot shows unique BCR counts (FR1) of primary tumors, retransplant and healthy donor cells. (b) Representative BCR network plots of deep sequenced PCR amplified immunoglobulin variable gene regions from primary tumor samples (n=8) and retransplants (n=4). Each vertex represents a unique sequence, where relative vertex size is proportional to the number of identical reads. Edges join vertices that differ by single nucleotide non-indel differences and clusters are collections of related, connected vertices. (c) Table shows selected mutations and their relationship in primary and retransplant tumors. (d) Unrooted phylogenetic trees of representative clonal expansions observed across samples. Each vertex is a unique BCR and branch lengths are estimated by maximum parsimony.

Overall, the ability of these tumors to closely recapitulate the appearances of high-grade B cell lymphoma further validates the biological relevance of this system to the study of human lymphoma and provides the opportunity to generate mutation-directed, bespoke *in vivo* lymphoma models.

## Discussion

The plethora of genomic information generated from next generation sequencing studies has left us with a need for new experimental systems in which to study the genetics of human lymphoma and to decipher these rich data resources. The availability and suitability of current preclinical models is recognized as a rate limiting step in translating genomic knowledge into patient benefit^11^. The cell of origin of most aggressive B cell lymphomas, including DLBCL and BL, is the GC B cell^12,13^. We therefore reasoned that non-malignant, human GC B cells should be the input for a system to create genetically defined models of human lymphoma. We describe an optimized system for the culture and transduction of primary, human GC B cells *ex vivo*. This relies on the provision of microenvironmental survival signals common to that of the germinal center, as well as the overexpression of combinations of oncogenes common to the pathogenesis of human lymphoma. In particular, this includes *BCL6*, a transcription factor central to the GC reaction as well as an established oncogene in GC-derived lymphoma. A related strategy has been employed previously to expand peripheral blood memory B cells for the purposes of monoclonal antibody engineering^40^. However, this is the first use of genetically altered human, primary, GC B cells for the functional investigation of lymphoma genetics and the first to generate synthetic, *in vivo*, human models of lymphoma.

A major advantage of using primary GC cells over established lymphoma cell lines is the ability to investigate defined genetic alterations on a genetically normal background. In particular, this provides a sensitive platform for investigating the ability of specific genetic alterations to increase survival and proliferation. An enhanced oncogenic phenotype is much harder to discern in cell lines where the mutational repertoire is likely to have evolved extensively for optimal *in vitro* growth. The superior sensitivity of this system, compared to cell lines, to detect alterations associated with increased growth or survival is evidenced by the strong enrichment for TSGs in our CRISPR screen when compared to conventional cell lines. The relevance of the system to the pathogenesis of human lymphoma is underscored firstly, by the ability to recapitulate the appearances of human high-grade B cell lymphoma *in vivo* and secondly, by the ability to identify GC-specific TSGs such as GNA13 in our CRISPR screen. Inactivating mutations of *GNA13* are common in lymphoma, but rarely seen in other forms of malignancy. Indeed, amplification is more common in solid organ cancers, where *GNA13* is generally considered to act as an oncogene^41^. The detection of *GNA13* missense mutations and a frameshift mutation of *S1PR2* in one of our synthetic lymphomas further underscores the importance of this pathway in GC derived lymphomas. We show how CRISPR/Cas9 can be integrated into the system for high throughput screening as well as for individual, gene-focused analysis such as the clear demonstration of AKT-independent survival advantage in *GNA13*-depleted cells. This is a finding consistent with the greater enrichment scores for *GNA13* compared to *PTEN* in our screening experiments.

The system affords versatility, with potential to vary the stimulation provided, the combination of expressed backbone oncogenes and the mechanism of their introduction. We envisage future studies that might remove or replace components of the feeder-based stimulation, for instance to identify factors promoting cytokine independent growth, or alter the background oncogene combination to screen for synergy between different oncogenic hits. The selective pressure imposed could be further altered by the use of pharmacological inhibitors of specific pathways. Future studies might also employ mutant open reading frame (mORF) screens or targeted CRISPR gene editing to introduce specific mutations into endogenous loci.

The complex genetic heterogeneity of human lymphoma is becoming increasingly evident^2–4^. It is clear that the repertoire of available cell lines does not adequately represent each of the many molecular subtypes predicted from the analysis of sequencing studies. Therefore, the ability to generate mutation-directed tumors *in vivo*, provides an attractive route for patient-personalized preclinical models. A particular advantage over tumor-derived xenograft models is the ability to create paired, syngeneic controls; tumors that are genetically identical other than the presence or absence of a specific mutation. Similar approaches to culture and manipulate human primary cells are proving successful for some solid organ malignancies^14–17^. However technical limitations have precluded this in B cell lymphoma. We present for the first time an extensively optimized, yet inexpensive strategy to employ primary, human, GC B cells for the investigation of lymphoma genomics and to generate bespoke, *in vivo* models of human lymphoma. This addresses an important bottleneck in translating lymphoma genomic findings into functional understanding that can drive improved patient outcomes and personalized therapy.

## Availability of data and reagents

Gene expression data has been uploaded to the EGA database under the accession number EGAS00001003560. All other remaining data are available within the article or supplementary information. Reagents including feeder cell lines, viral packaging line, viral plasmids will be distributed freely upon request.

## Acknowledgements

D.H. was personally supported by a Clinician Scientist Fellowship from the Medical Research Council (MR/M008584/1). Research in the Hodson laboratory is supported by the Kay Kendall Leukaemia Fund, The Addenbrooke’s Charitable Trust, Cancer Research UK and a research scholarship from Gilead Sciences. The laboratory receives core funding from Wellcome and MRC to the Wellcome-MRC Cambridge Stem Cell Institute. We thank Alice Mitchell and the ENT Department at Addenbrooke’s Hospital, Cambridge for their assistance in the collection of primary tonsil tissue. We are grateful to Joanna Baxter and Cambridge Blood and Stem Cell Bank for collection and storage of primary tonsils samples and to the staff of the Central Biomedical Services for animal housing and care. We thank Craig McDonald for technical assistance with video microscopy and Martin Dyer, Ingo Ringshausen and Reuben Tooze for critical reading of the manuscript.

---

**Figure S1.**
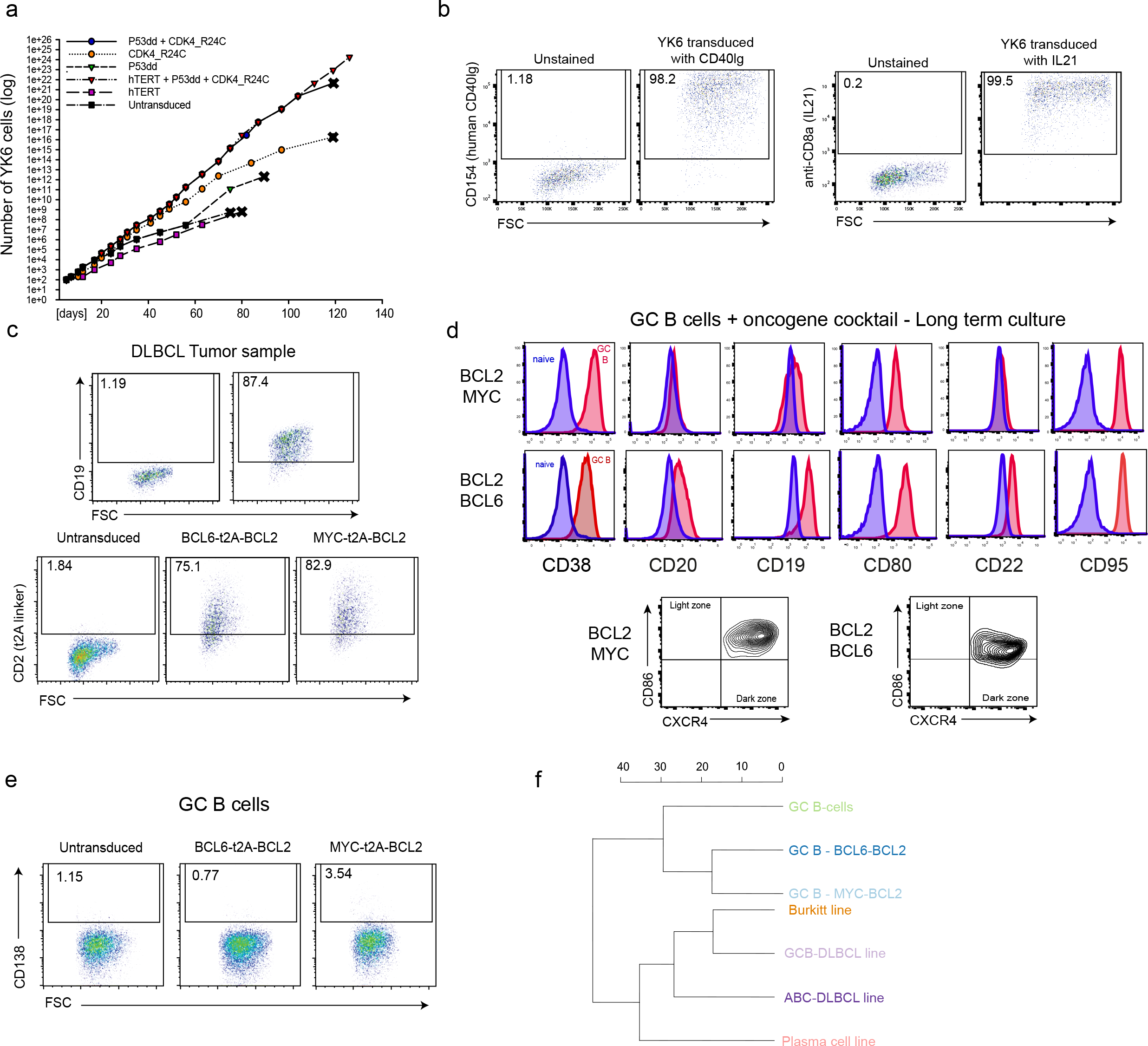
(a) Immortalization of the FDC-like feeder cells (YK6) was achieved by retroviral transduction of the indicated oncogenes. Cell number was monitored by manual cell counting. Illustrated is time course showing the number of cells following transduction. Cohorts that had stopped proliferating and underwent replicative senescence are marked with an × at their final timepoint. (b) Flow cytometry analysis for the expression of CD40lg (CD154) and IL21 (transduction marker CD8a) on immortalized FDC-like cells that were engineered to express CD40lg and IL21. FSC, forward scatter. (c) DLBCL primary tumor sample was transduced with retroviral vectors expressing *BCL2*-t2A-*BCL6* and *BCL2*-t2A-*MYC* using the MuLV-GaLV fusion envelope. Cells were stained for transduction marker CD2 (t2A) and B cell marker CD19 and were analyzed by flow cytometry. (d) Primary human GC B cells were transduced with the oncogenic cocktail *BCL2-BCL6* and *BCL2-MYC* and cultured long term (Day 73). Representative flow cytometry analysis (*n* = 3) for the expression of the GC B cell markers CD38, CD20, CD19, CD80, CD22, CD95, CXCR4 and CD86. Red histograms show GC B cells compared to primary human naïve B cells (blue). (e) Flow cytometry analysis for the expression of CD138 in cultured primary human GC B cells transduced with *BCL2-BCL6* or *BCL2-MYC* was performed 4 weeks after transduction. FSC, forward scatter. (f) Subtypes of cells in Figure 2E were compared using a signature of germinal centre expressed genes (GCB-1). The clustering was based on the Euclidean distance between the average normalized expression.

**Figure S2.**
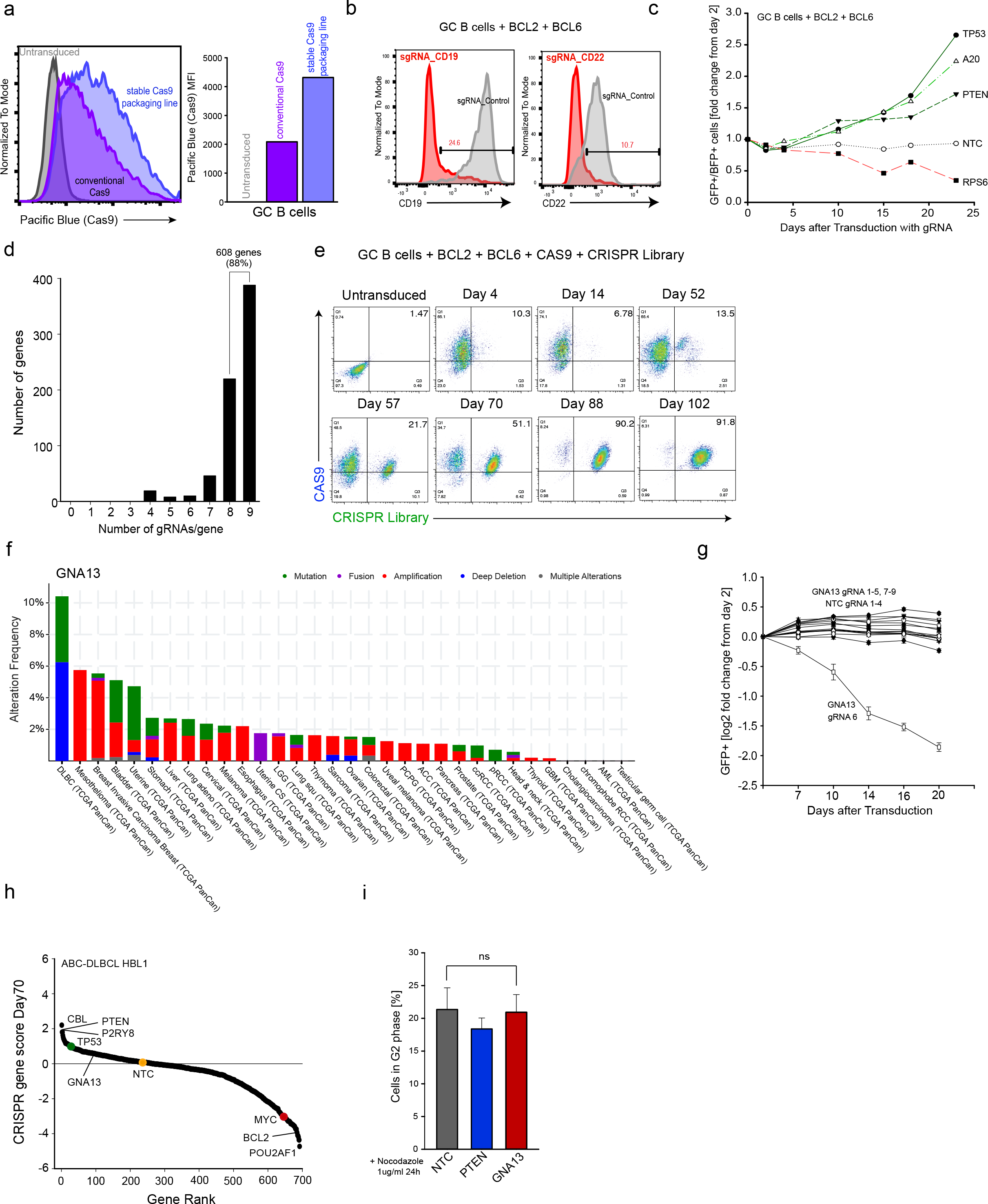
(a) Two different methods of Cas9 virus production were compared; conventional transfection of 293T packaging cells with retroviral Cas9 construct, and a stable Cas9 packaging line, both tagged with BFP. Bar chart illustrates the mean fluorescence intensity of Pacific Blue (Cas9) in primary GC B cells transduced with each method of Cas9 virus production. (b) Primary GC B cells were transduced with *BCL2-BCL6* and Cas9-BFP and subsequently with gRNAs against *CD19, CD22* and non-targeting control. Staining for CD19 and CD22 was performed 6 days after gRNA transduction and gated on double positive CAS9 (BFP) and gRNA (GFP) expressing cells. Red histograms show CD19/CD22 expression in cells transduced with the indicated gRNA. Grey histograms show expression of CD19/CD22 transduced with a non-targeting control. (c) Primary GC B cells were transduced with *BCL2-BCL6* and Cas9-BFP and subsequently with gRNAs against *TP53, PTEN, A20, RPS6* and non-targeting control (NTC). Following transduction with gRNAs, enrichment or depletion of BFP^+^GFP^+^ cells was monitored by flow cytometry. Illustrated is time course showing the relative changes of BFP^+^GFP^+^ cells following transduction. (d) Illumina sequencing revealed the number of gRNAs present per gene in the lymphoma-focused CRISPR library. (e) Flow cytometry analysis showing enrichment of double positive cells transduced by both Cas9-BFP and CRISPR library (GFP) over different timepoints in primary human GC B cells transduced with *BCL2-BCL6*. (f) TCGA analysis of *GNA13* alteration frequency in different cancer types. Figure obtained from cBioPortal^34,35^. (g) HBL1-Cas9 cells were transduced with 9 *GNA13*, 4 *PTEN* and 4 non-targeting control gRNAs and enrichment or depletion of GFP^+^ cells was monitored by flow cytometry. Illustrated is time course showing the log2 fold-change relative to baseline (± Standard Deviation) of GFP^+^ cells following transduction (n = 3). Representative of 3 experiments with 3 replicates/experiment. (h) HBL1-Cas9 was transduced with the lymphoma-focused CRISPR library. Genes are ranked from highest to lowest according to their CRISPR gene scores at Day 70 (log2 scale). Selected tumor suppressor genes as well as oncogenes are highlighted in green and red, respectively. Everything above the horizontal line is positively enriched. (i) Cell Proliferation following *GNA13* and *PTEN* deletion in primary human GC B cells was monitored by Vybrant™ DyeCycle™ Ruby Stain and analyzed by flow cytometry (n=3). GC B cells were treated with Nocodazole (1ug/ml) 24h prior to FACS analysis to arrest cells in G2 phase. Bar chart illustrates G2 positive cells.

**Figure S3.**
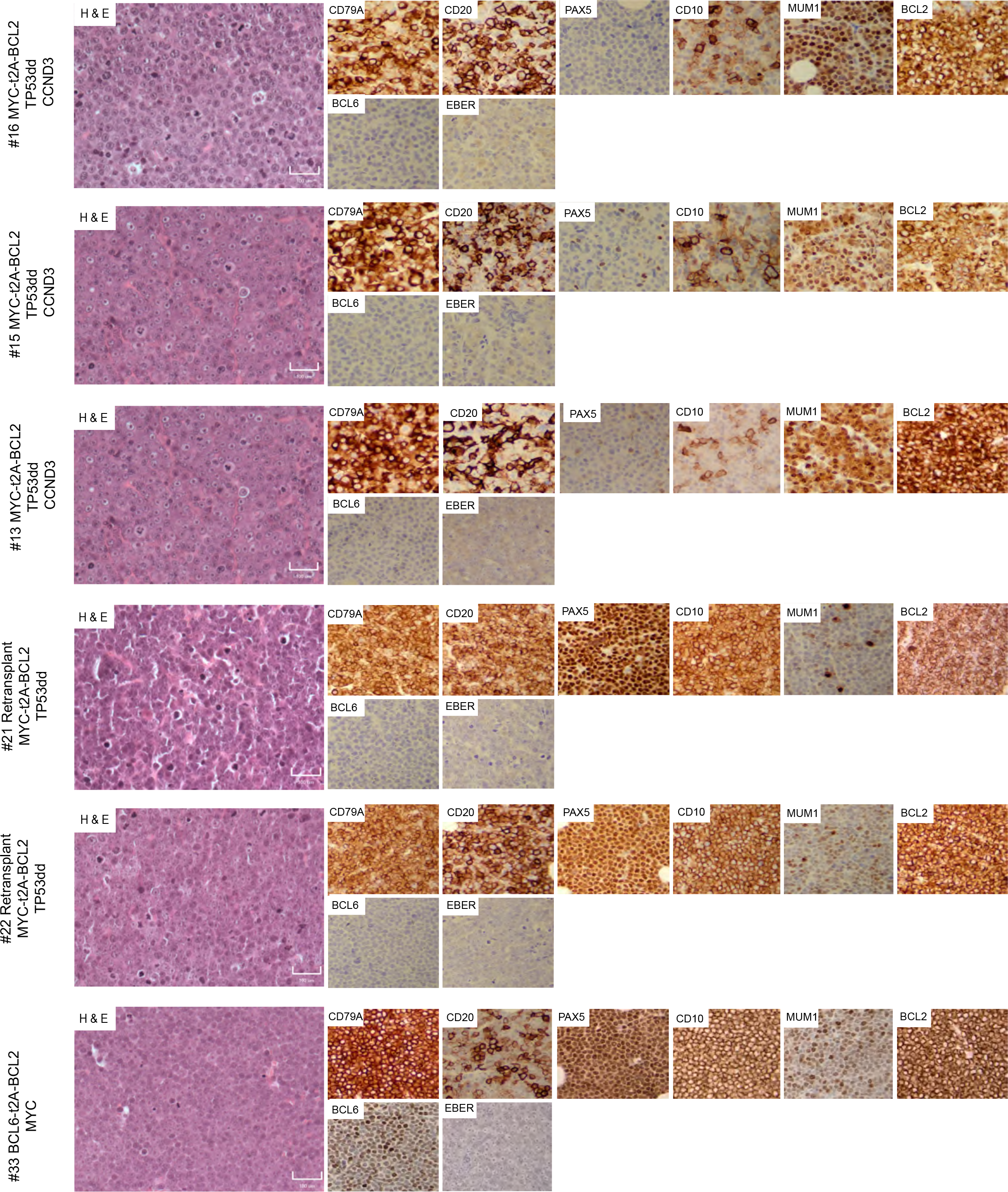
Immunohistochemistry images for the indicated markers are shown (Magnification 20×). Six different tumors are shown. Scale bar, 100μM.

**Figure S4.**
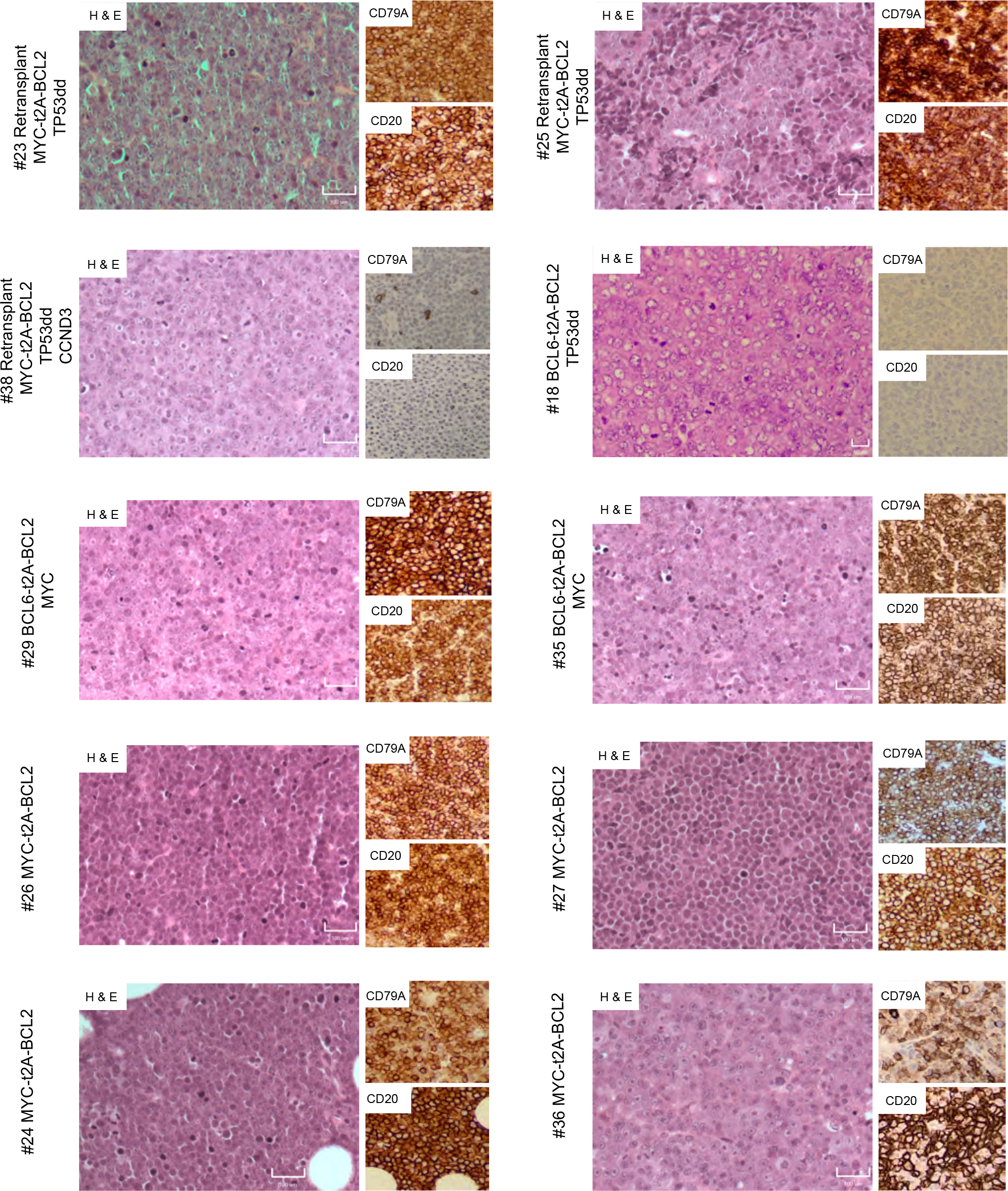
Immunohistochemistry images for H&E, CD79A and CD20 are shown (Magnification 20×). Ten different tumors are shown. Scale bar, 100μM.

**Figure S5.**
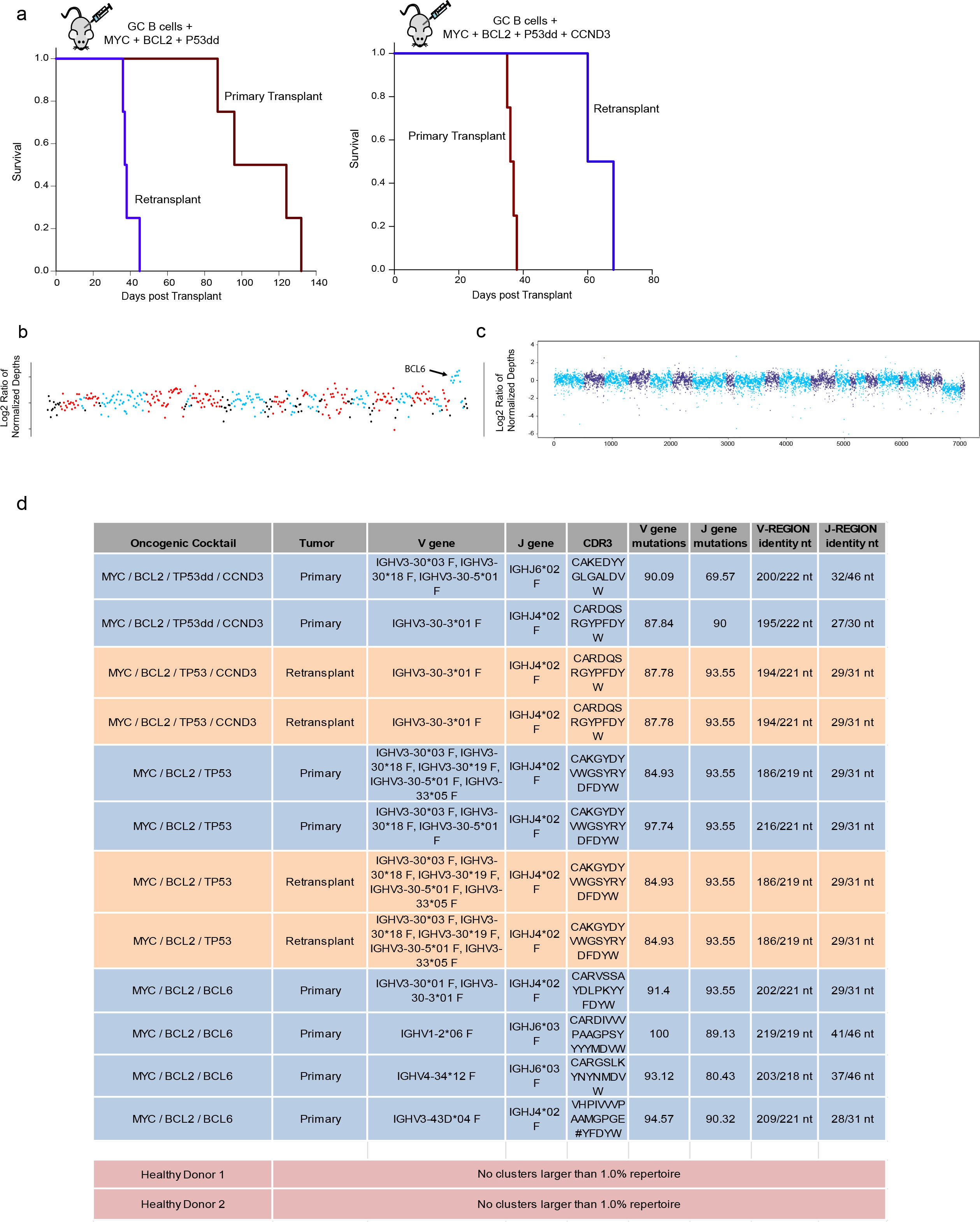
(a) Primary human GC B cells harvested from primary tumors were retransplanted subcutaneously into NOD/SCID/gamma mice and monitored for palpable tumors. Mice were culled when tumors reached 12mm in size. Overall survival of the recipient mice (Primary transplant and retransplant) is plotted as a Kaplan-Meier curve. (b) Gains corresponding to BCL6 on chr3 are shown. Black points are background regions whilst red and blue features correspond to genes analyzed for their copy number state. BCL6 is the last blue feature on the chromosome. (c) Copy number across whole genome showing no evidence of aneuploidy. Alternating colors correspond to different chromosomes. The x-axis represents bin index rather than absolute genomic coordinate. (d) Table for V and J segment mutations and identity (nt) shown for primary tumor samples (n=8) and retransplants (n=4).

***Supplementary Videos***

Videos show culture of GC B cells alone, YK6 alone, YK6 + GC B cells, YK6-CD40Lg + GC B cells and YK6-CD40Lg-IL21 + GC B cells from the time of plating to 132 hours after. Scale bar, 50μM.

***Supplementary Table 1***

Enrichment scores based on relative read counts of barcoded expression constructs for transcription factors or their mutant versions in GC B cells co-transduced with *BCL2* over 4 different timepoints (n=3).

***Supplementary Table 2***

CRISPR gene scores shown for GC B cells transduced with *BCL2*, *BCL6* (n=3) and cell line HBL1 (n=1) with the CRISPR library targeting 692 genes (n=3).

***Supplementary Table 3***

Protein altering variants identified by MUTEC2 using the matched, pre-transduced, germinal center B cells as the normal control.

## Online Methods

### Plasmid Construction

The CDS for human gene sequences were cloned into the pBMN-IRES-LyT2 retroviral vector (kind gift of Dr Louis Staudt, National Cancer Institute, USA)^1^ to express *CCND3*, *BCL6* or *IL21* using synthetic double-stranded DNA from IDT with Gibson Assembly (NEB). For *CCND3*, a mutation in the threonine residue at Threonine 283 (T283A) was included, to enhance protein expression^2^. *BCL2, CD40L* or *MYC* were cloned into MSCV-based vectors using synthetic double-stranded DNA and Gibson Assembly. For overexpression of multiple human genes, CDS sequences were cloned into the MSCV-IRES-huDC2 vector (kind gift of Dr Martin Turner, the Babraham Institute, UK) with the t2A peptides linking genes, such as BCL6-t2A-BCL2, MYC-t2A-BCL2 and P53dd-t2A-BCL2 or an additional p2A peptide when linking three genes such as P53dd-p2A-CDK4_R24C-t2A-BCL2. pBABE-hygro-hTERT was a gift from Bob Weinberg (Addgene plasmid # 1773; http://n2t.net/addgene:1773; RRID:Addgene_1773)^3^. P53dd and CDK4_R24C were cloned into MSCV-IRES-Thy1.1 DEST vector using synthetic double-stranded DNA from IDT with Gibson Assembly. MSCV-IRES-Thy1.1 DEST was a gift from Anjana Rao (Addgene plasmid # 17442; http://n2t.net/addgene:17442; RRID:Addgene_17442)^4^.

The MSCV-CAS9-2A-BFP construct was modified from Addgene plasmid #65655 by excision of the IRES/Puro elements and insertion of a 2A-BFP sequence using dsDNA and Gibson assembly. MSCV_Cas9_puro was a gift from Christopher Vakoc (Addgene plasmid # 65655; http://n2t.net/addgene:65655; RRID:Addgene_65655)^5^.

The custom CRISPR library and single gRNAs were cloned into pKLV2-U6gRNA-Bbsi-PGK-GFP, which was modified from pKLV2-U6gRNA5(Empty)-PGKBFP2AGFP-W. pKLV2-U6gRNA5(Empty)-PGKBFP2AGFP-W was a gift from Kosuke Yusa (Addgene plasmid # 67979; http://n2t.net/addgene:67979; RRID:Addgene_67979)^6^. Pooled oligos for construction of the lymphoma-focused CRISPR library were obtained from TWIST Bioscience and oligos for single gRNAs were obtained from IDT. To make the GaLV-MuLV fusion envelope constructs, pHIT123^7^ (kind gift of Prof Markus Muschen, City of Hope, Los Angeles, CA) containing the retroviral ecotropic envelope, human cytomegalovirus immediate-early promoter and the origin of replication from simian virus 40 was used as the backbone. The viral envelopes GaLV_WT, GaLV_MTR and GaLV_TR were based on the SEATO strain of GaLV (NP_056791). GaLV_MTR and GaLV_TR contain the 3’ GaLV envelope sequence replaced by the MuLV transmembrane region, cytoplasmic region and R peptide region and the MuLV cytoplasmic region and R peptide region, respectively^8^. All sequences were purchased from IDT as synthetic double-stranded DNA and inserted by Gibson assembly. All plasmids were verified by capillary sequencing.

### Cell culture

Cell lines HBL1, BJAB, U2932, TMD8, SUDHL4, DOHH2, Raji, Mutu (all kind gifts from Dr Louis Staudt, National Cancer Institute, USA) and NCIH929 (kind gift from Dr Mike Chapman, University of Cambridge) were cultured in Roswell Park Memorial Institute medium (RPMI-1640, Invitrogen, Carlsbad, CA). Primary human GC B cells were cultured in Advanced Roswell Park Memorial Institute medium (Advanced RPMI-1640, Invitrogen, Carlsbad, CA) with GlutaMAX containing 20% FBS, 100 IU/ml penicillin and 100 µg/ml streptomycin and kept at 37°C in a humidified incubator (5% CO_2_ and 95% atmosphere). Lenti-X 293T Cell Line (Clontech Laboratories) were cultured in Dulbecco’s Modified Eagle Medium (DMEM, Invitrogen, Carlsbad, CA) containing 10% FBS, 100 IU/ml penicillin and 100 µg/ml streptomycin and kept at 37°C in a humidified incubator (5% CO_2_ and 95% atmosphere). All cell lines used in this study were confirmed to be free from mycoplasma contamination and identity was verified using a 16-amplicon multiplexed copy number variant fingerprinting assay^9^.

### Construction of YK6-CD40Lg-IL21 feeder line

Discarded human tonsil tissue was obtained after a routine tonsillectomy and handled in accordance with an IRB-approved protocol (2013-0864) at the Asian Medical Center, Seoul, South Korea. Follicular dendritic cells were extracted from tonsils following an established protocol for the creation of HK FDC-like feeder cells^10^. Following mechanical disruption and enzymatic digestion, the released cells were collected and subjected to Ficoll gradient centrifugation for 20 min at 2,200 rpm. The interface layer that contains FDC was then collected. The cells were resuspended in RPMI 1640 medium and centrifuged at 200 rpm for 10 min at 4°C over a discontinuous gradient of 7.5% and 3% bovine serum albumin (BSA; A9418, Sigma-Aldrich, St. Louis, MO, US). FDC-enriched fractions were collected from the interface. Cells were washed with HBSS and cultured on tissue culture dishes. Cells isolated and culture after these procedures initially contained large adherent cells with attached lymphocytes. Non-adherent cells were removed and adherent cells replenished with fresh medium every 3-4 days. Adherent cells were trypsinized when confluence was attained. Because of the limited growth in culture, FDC-like cells were immortalized (now termed YK6) through retroviral transduction with pBABE_TERT.Hygro, P53DD_Thy1.1 and CDK4_R24C_Thy1.1. Immortalized YK6 cells were further transduced with hCD40Lg-Puro and IL21-LyT2.

### Purification of human Germinal Center B cells

Fresh, tonsil tissue was sourced from the Addenbrooke’s ENT Department, Cambridge and processed directly to preserve viability. Ethical approval for the use of human tissue was granted by the Health Research Authority Cambridgeshire Research Ethics Committee (REC no. 07/MRE05/44). Germinal center B cells were purified using the human B cell negative selection isolation Kit II (MACS, Miltenyi Biotec) as per manufacturer’s instructions. The protocol was modified to include negative selection antibodies IgD-BIOT (SouthernBiotech) and CD44-BIOT (SouthernBiotech) to remove naïve and memory B cells^11^. Cells were stained for CD38, CD20, CD19 and CD10 (Biolegend) to confirm enrichment of germinal center B cells. B cell genomic DNA was screened for EBV status using a quantitative real-time PCR (qPCR) assay^12^ and cells from EBV-positive were discarded. GC B cells were plated onto irradiated YK6-CD40lg-IL21 and split every 2-3 days. Fresh YK6-CD40lg-IL21 cells (irradiated 30 Gy) were added with each split. Primary human GC B cells were cultured in Advanced Roswell Park Memorial Institute medium (Advanced RPMI-1640, Invitrogen, Carlsbad, CA) with GlutaMAX containing 20% FBS, 100 IU/ml penicillin and 100 µg/ml streptomycin and kept at 37°C in a humidified incubator (5% CO_2_ and 95% atmosphere).

### Retroviral and lentiviral production

Retroviral packaging plasmids pHIT60 (kind gift of Dr Louis Staudt, National Cancer Institute, USA) and GaLV WT were used as follows: 1μg pHIT60 (gag-pol), 1μg GaLV WT (envelope) and 4μg of a retroviral construct was used to transfect each 10 cm^2^ dish of HEK-293T, after mixing with 1 ml of Opti-MEM media (Invitrogen) and 18μl of TransIT-293 (Mirus). For lentivirus transfections, packaging plasmids pCMVDeltaR8.91 and GaLV MTR were used as follows: 8.3 μg pCMVDeltaR8.91 (gag-pol), 2.8μg GaLV MTR (envelope) and 11μg of a lentiviral construct per 10 cm^2^ dish, incubated with 1 ml of Opti-MEM media (Invitrogen) and 33μl of TransIT-293 (Mirus). For infecting cell lines, pMD2.G (VSV-G envelope) was used instead of GaLV MTR. pMD2.G was a gift from Didier Trono (Addgene plasmid # 12259; http://n2t.net/addgene:12259; RRID:Addgene_12259). The packaging line Lenti-X 293T Cell Line (Clontech Laboratories) was used for all transfections. After 36-48h, the virus supernatant was filtered through a 0.45μM filter. If needed, media was replenished and harvested again at 16 hours later. For retroviral/lentiviral transduction, 1-2 × 10^6 target cells were resuspended with viral supernatant and infected by centrifugation (1500 × g, 90 min at 32°C) with the addition of 10μg/ml Polybrene (INSIGHT biotechnology) and 25μM HEPES (ThermoFisher) in 12 or 24 well plates. Viral supernatant was replaced with fresh media immediately after centrifugation for retroviral infection or after > 4 hours if transducing lentiviral constructs. Cells were maintained at 37°C with 5% CO2 for 2 - 4 days before analyzing by FACS.

### Mouse experiments

Cultured human germinal center B cells were injected subcutaneously into the left flank of NSG mice (Jackson laboratories). Up to 10 × 10^6 cells were washed and resuspended with Matrigel (Corning) in a 1:1 ratio. Mice were culled when tumors reached 12 mm in size. Tumors were processed immediately after harvest and analyzed by flow cytometry and histology. This research has been regulated under the Animals (Scientific Procedures) Act 1986 Amendment Regulations 2012 following ethical review by the University of Cambridge Animal Welfare and Ethical Review Body (AWERB).

### Generation of lymphoma-focused CRISPR guideRNA library

gRNA sequences were based upon two recent genome-wide libraries^6,13^. Position one of 20 set to G for all gRNAs. Appropriate overlapping sequences (underlined) for Gibson Assembly into the gRNA expression plasmid pKLV2_U6gRNA_Bbsi_PGK_GFP (modified from Addgene #67979) were appended to all 6000 gRNAs. A 70-mer oligo pool was purchased from TWIST BIOSCIENCE as follows: 5’-<u>TATCTTGTGGAAAGGACGAAACACCG-</u>N_19_- <u>GTTTAAGAGCTATGCTGGAAACAGC</u>-3’ N_19_ represents each of the 6000 gRNA sequences. The single-stranded oligo pool was converted to double-stranded DNA by PCR amplification using Q5 Hot Start High-Fidelity 2X Master Mix (NEB) with 3 ng of the oligo pool as a template and primers (Zhang_F and Zhang_R_modified). The following PCR conditions were used: 95 °C for 2 min, 10 cycles of 95 °C for 20 s, 60 °C for 20 s and 72 °C for 30 s, and the final extension, 72 °C for 3 min. The 150bp PCR product was gel purified from a 2% Agarose gel using the Gel Extraction kit (Qiagen) and eluted in 20μl EB Buffer. Four Gibson Assembly reactions were performed using 14.4 ng of the purified 150bp fragment and 200 ng of the BbsI-digested pKLV2_U6gRNA_Bbsi_PGK_GFP with Gibson HiFi DNA Assembly Master Mix (NEB). Gibson Assembly reactions were pooled and column-purified using MinElute PCR purification kit (QIAGEN). Eight electroporations were performed using 1μl of the purified Gibson reaction and 20μl of Endura Competent Cells (Lucigen). The mixture was transferred to a 0.1 cm cuvette and electroporated at 1.8KV. Immediately after, 2 ml of prewarmed SOC media was added to each reaction and placed on a shaker at 37°C for one hour. The electroporated cells were combined and plated onto sixteen 24.5 cm^2^ LB + ampicillin agar plates using ColiRollers Plating Beads (Merck Millipore). Plates were left at 30°C overnight and plasmid DNA was purified using a Plasmid Maxi kit (Qiagen).

### Transduction of CRISPR library and generation of gRNA sequencing libraries

GC B cells were transduced with the backbone oncogene cocktail and Cas9-BFP retrovirus until Cas9-BFP reached between 50 and 80%. The number of cells transduced with gRNA library was adjusted to take account of the percentage of Cas9-expressing cells and target MOI of 0.3 in order to maintain representation of >1000× the size of the library. Four days after transduction, BFP and GFP expression was analyzed by flow cytometry and at each harvest timepoint going forward. A minimum of 1000× representation was maintained at each passaging step. Cells were harvested every 14 days. Genomic DNA extraction was conducted as described previously^14^ and Illumina sequencing was performed as described^6,15^. Purified libraries were quantified, pooled and sequenced on Illumina HiSeq4000 by 50-bp single-end sequencing.

### Computational analysis of CRISPR screens

Raw reads were normalized to a total number of reads in a sample as follows:

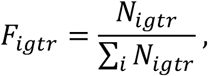

denotes the raw sequencing reads of gRNA *i* of gene *g* at time *t* in replicate *r*. For each gRNA the Z-score of log_2_ fold change between plasmid library and late sample, *Z*_*igr*_ is given by:

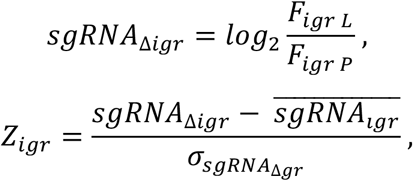

Finally, *CRISPR score*_*g*_, which represents the magnitude and direction of a fitness of a gene *g* between the two time points is:

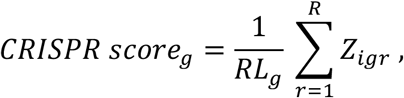

where *L*_*g*_ denotes the number of sgRNA of gene *g* in replicate *r* and *R* is the number of available replicates.

### RNA-Sequencing

Total RNA from cells was extracted using NucleoSPIN RNA from Macherey-Nagel and cDNA was produced from 500ng of total RNA using qScript™ cDNA SuperMix (Quanta Biosciences). RNA-seq library was prepared using the NEBNext Poly (A) mRNA Magnetic Isolation Module (E7490) and NEBNext Ultra Directional RNA Library Prep Kit for Illumina (E7420) according to manufacturer’s instructions. NEBNext Multiplex Oligos for Illumina (E7500) was used for indexing and sequenced on a HiSeq4000 by 50-bp single-end sequencing. RNA-Seq were mapped to the hg38/GRCh38 reference human genome using splice-aware aligner STAR 2.5.3a^1^ in two pass-mode. The genome index was built with GENCODE v.28 comprehensive gene annotation set. Uniquely mapped reads were assigned to genes with Rsubread package^2^ allowing for assignment of a read to more than one overlapping features. At least 25 of overlapped bases were required to assign a read to a gene. Genes with low-counts were filtered out with a threshold of minimum 128 counts in at least 25% of samples. Gene expression values were obtained using variance stabilizing transformation as implemented in the DESeq2^3^ package.

### Barcoded overexpression experiments

The CDS for human gene sequences (BCL6 WT, BCL6_G559R, BCL6_H641R, BCL6_R585W, BCL6_P586A, IRF8 WT, IRF8_N87Y, IRF8_T80A, IRF8_380_stop, IRF8_S55A, MEF2B WT, MEF2B_D83V, MEF2b Y69H) were cloned into barcoded pBMN-IRES-LyT2 retroviral vector using NEBuilder^®^ HiFi DNA Assembly. Primary human GC B cells were retrovirally transduced with BCL2, followed by infection with barcoded overexpression genes and pooled four days after transduction, and then grown in competitive culture. Genomic DNA was collected at day 4 and approximately every 14 days after. Genomic DNA extraction was conducted as described previously^14^ and individual indices (Truseq Small RNA Index Sequences) were added using Q5 Hot Start High-Fidelity 2× Master Mix. The purified library was quantified, pooled and sequenced on Illumina MiSeq by 50-bp single-end sequencing.

### Computational analysis of barcoded overexpression experiments

Relative abundances *F*_*ictr*_ of a construct in a pooled competitive culture were computed as follows:

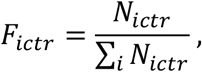

where *N*_*ictr*_ denotes the raw sequencing counts of clone *i* of constructs *c* at time *t* in replicate *r*. The average relative abundance of construct *c* at time *t* is given by:

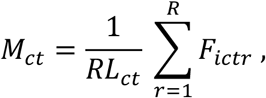

where *L*_*ct*_ denotes the number of clones of construct *c* at time *t* and *R* is the number of available replicates.

### BCR amplification

PCR amplification of DNA from synthetic lymphoma tumors (100 ng input) was performed with JH reverse primer and FR1 forward primer set pools (provided by Sigma Aldrich) as previously described^16^. MiSeq libraries were generated using KAPA Hyper Prep Kit (KAPA Biosystems) incorporating KAPA Dual-Indexed Adapter for Illumina MiSeq platforms and reads were filtered as described previously^17^. The computational pipeline MRD Assessment and Retrieval Code in Python (MRDARCY) was then used to analyze BCRs, followed by secondary rearrangement analysis in which the relative frequencies of each IgHV gene were determined by BLAST using the IMGT reference gene database^18^. The following primers were used:

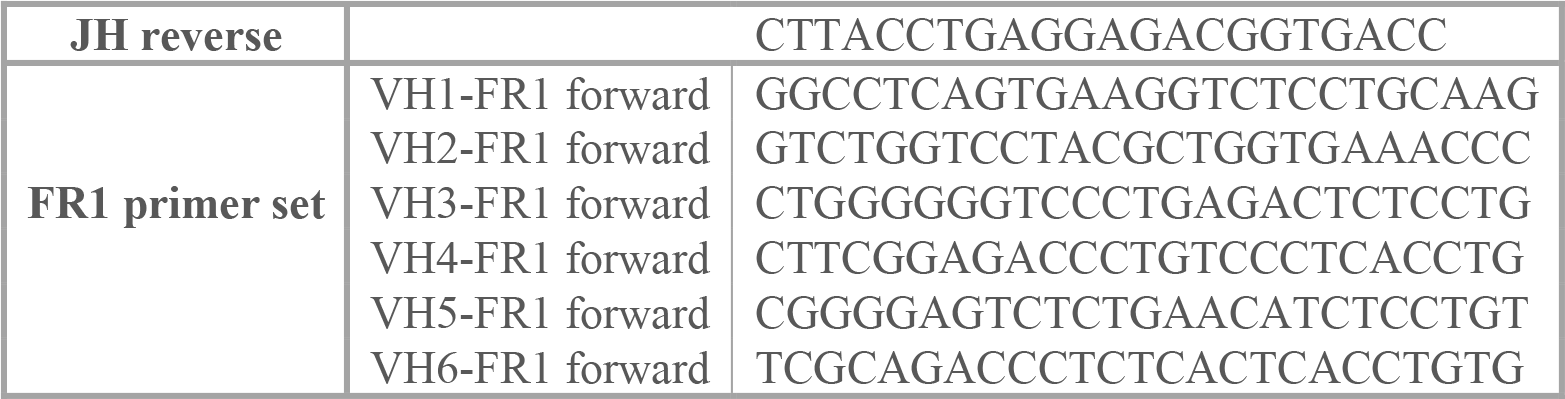

### High-throughput sequencing and analysis of heavy chain immunoglobulin

Deep sequencing of PCR amplified immunoglobulin heavy chain variable gene regions and BCR network generation algorithm and network properties were performed as previously described^16^ Each vertex represents a unique sequence, where relative vertex size is proportional to the number of identical reads. Edges join vertices that differ by single nucleotide non-indel differences and clusters are collections of related, connected vertices. Ig gene usages and sequence annotation were performed in IMGT V-QUEST, where repertoire differences were performed by custom scripts in Python.

For the visual representations of the BCR repertoires, BCR network subsampling was performed using the cluster-enforced linkage sampling (CC) method to preserve the overall clonal structure. Briefly, the CC algorithm employs three steps to account for loss of connectivity between vertices in clusters during sampling:

1. *Vertex selection:* Vertices were reselected until the number of desired clusters in the original network G are represented.
2. *Cluster-vertex migration:* For each cluster in the original network which contains more than one vertex that was sampled, vertices were reselected such that the cluster connectivity is retained in the sampled network.
3. *Induced graph formation:* Graph induction selects the set of edges (Es) to be included in the sampled graph. Total graph induction is used in CC, selecting all edges incident on the sampled vertices are included in the sampled graph.

This process was repeated 20 times, and the subsample that most closely represented the true (unsampled) maximum cluster size was retained and plotted IGHV gene editing analyses were performed in a similar manner to^18^. For all BCRs, stem regions were identified (defined as N-IgHD-N-IgHJ regions starting 3bp downstream of the IgHV gene boundary). The number of unique BCR sequences sharing stem regions but with different IgHV gene usage (>95% difference in sequence identity in the IgHV region) and with different 5’ of the junctional region (defined as IgHV(last 3pb)-N-IgHD-N-IgHJ) was determined and compared to the total number of unique BCRs to give the percentage *IgHV* replacement. Sequences with joining regions (N-IgHD-N-IgHJ regions) shorter than 8 nucleotides were excluded from this percentage due to potential of germline encoded receptors.

### Mutation analysis

To identify somatic mutations across synthetic lymphoma tumors a hybrid-capture platform was used with a bait set^14^ (SureSelect, Agilent, UK, ELID # 0731661) of 292 genes frequently mutated in hematological malignancies. After hybridization-based sequence enrichment (SureSelect^HSXT^, Agilent), high-throughput sequencing was performed on the Illumina HiSeq 4000 platform.

### Sequencing data alignment

DNA sequencing reads were aligned to the GRCh37d5 according to the workflow described at Samtools webpage for mapping and improvement (http://www.htslib.org/workflow/) using BWA-MEM^19^ 0.7.17, Samtools^20^ 1.9, GATK^21^ 3.8.1 and Picard (http://broadinstitute.github.io/picard/) 2.18.25.

### Variant calling for substitutions and indels

Single base substitutions and short insertions and deletions were called using GATK^21^ 4.1 Mutect2 based on the tutorials available at Broad Institute website (https://gatkforums.broadinstitute.org/gatk/discussion/11136/how-to-call-somatic-mutations-using-gatk4-mutect2). The mutant variants were annotated using Variant Effect Predictor^22^ from ENSEMBL version 95.

### Copy Number Analysis

Copy number analysis was performed using GeneCN (https://github.com/wwcrc/geneCN).

### FACS (Fluorescence-activated cell sorting)

Cells were stained with Fluorophore-labelled antibodies in 2% BSA in PBS according to manufacturer’s instructions. The stained/or unstained cells were analyzed on the LSRII (BD). For cell counting, CountBright Absolute Counting Beads (ThermoFisher) were used according to manufacturer’s instructions and analyzed on the LSRII (BD). For dead cell apoptosis analysis, APC-conjugated Annexin V/Dead Cell Apoptosis Kit (Life Technologies) was used for the detection of apoptotic cells according to the manufacturer’s instructions. Externalization of phosphatidylserine (Annexin V, APC Conjugate) and DNA content (7-AAD) were measured and gating on all cells was used for further analysis. Cell cycle analysis was performed using the Vybrant^®^ DyeCycle™ Ruby Stain (Thermo) according to manufacturer’s instructions. Cells were treated with Nocodazole (1ug/ml) 24h prior to staining. Stained cells were analyzed by gating on cells in G2 phase using FlowJo software. Intracellular staining of phosphorylated AKT was performed as follows: Cell suspension and pre-warmed Fixation Buffer (BD Cytofix) was gently mixed in a 1:1 ratio and incubated at 37°C for 15min. Cell suspension was pelleted and washed with PBS twice at 350g for 5 min. Ice-cold True-Phos perm buffer (BD Cytofix) was added dropwise to the cell pellet whilst vortexing, followed by incubation at −20°C for at least 60 min. Cells were further washed twice and resuspended in FACS buffer (PBS + 2% FBS) containing the appropriate antibody at a dilution of 1:50 (Phospho-Akt Ser473, Cell Signaling, #11962). After staining for 30min, cells were washed and resuspended in FACS buffer followed by analysis on the LSRII (BD). The following antibodies were used: CD38 (HB7), CD20 (2H7), CD19 (HIB19), CD10 (97C5), CD2 (RPA-2.10), CD90.1 Thy1.1 (OX-7), CD154 (24-31), CD8a (53-6.7), CD22 (HIB22), CD80 (2D10), CD40 (5C3), CD95 (DX2), CD86 (IT2.2) and CXCR4 (12G5). All antibodies were purchased from Biolegend.

### Western blotting

Western blotting was performed as described previously^14^. The following antibodies were used: Beta-actin (13E5, Cell Signaling Technology) and GNA13 (EPR5436, Abcam).

